# Compound design of a patient-derived 3D cell culture system modelling early peritoneal endometriosis

**DOI:** 10.1101/2025.03.17.643554

**Authors:** Muhammad Dimas Reza Rahmana, Christopher J. Hill, Bettina Wilm, Dharani K. Hapangama

**Author notes:** Joint senior authors (Bettina Wilm, Dharani K. Hapangama).

## Abstract

Peritoneal endometriosis causes pelvic pain and infertility, but the underlying mechanisms related to these symptoms are not fully understood. Endometriosis diagnosis is typically delayed; thus, patient samples are unsuitable to study early endometriosis formation *in situ*. This study aimed to generate a 3D co-culture model of early peritoneal endometriosis using patient-derived primary cells, providing unique opportunities to examine endometriotic lesion initiation and progression. Peritoneal wash fluid, fallopian tube mesentery and endometrial biopsies were collected from patients undergoing laparoscopic surgery to isolate primary cells. A composite 3D hydrogel construct was assembled by embedding human peritoneal fibroblasts (HPFs) in a Matrigel-collagen I matrix and subsequent seeding with a layer of human peritoneal mesothelial cells (HPMCs). Immunohistological investigation of the composite hydrogel construct confirmed the successful assembly of a simple peritoneum layer model comprising a mesothelial monolayer, basement membrane and underlying fibroblasts, while secretion of tissue plasminogen activator demonstrated functional mesothelial physiology. Endometrial epithelial organoids (EEOs) were co-cultured with endometrial stromal cells (ESCs) to form endometrial assembloids mimicking shed endometrial tissue fragments at menstruation. When transplanted onto the peritoneal layer model, endometrial assembloids adhered, thus simulating early endometriotic lesion formation. Histological analysis demonstrated direct cell-cell contacts between HPMC, HPF and ESC at the endometrial-peritoneal interface, suggesting the involvement of those cell types in lesion initiation. Our modifiable superficial endometriosis model allows for further refinement by the addition of hormones, cytokines and/or other cell types to determine the underlying molecular mechanism involved in endometriotic lesion formation.

**Highlight:**

- Simple peritoneal layer model resembles parietal peritoneum structure and function.
- Combined endometrial and peritoneal model mimics early endometriosis formation.
- The patient-derived multi-cellular model is suitable to study early endometriosis.

**Graphical abstract:** 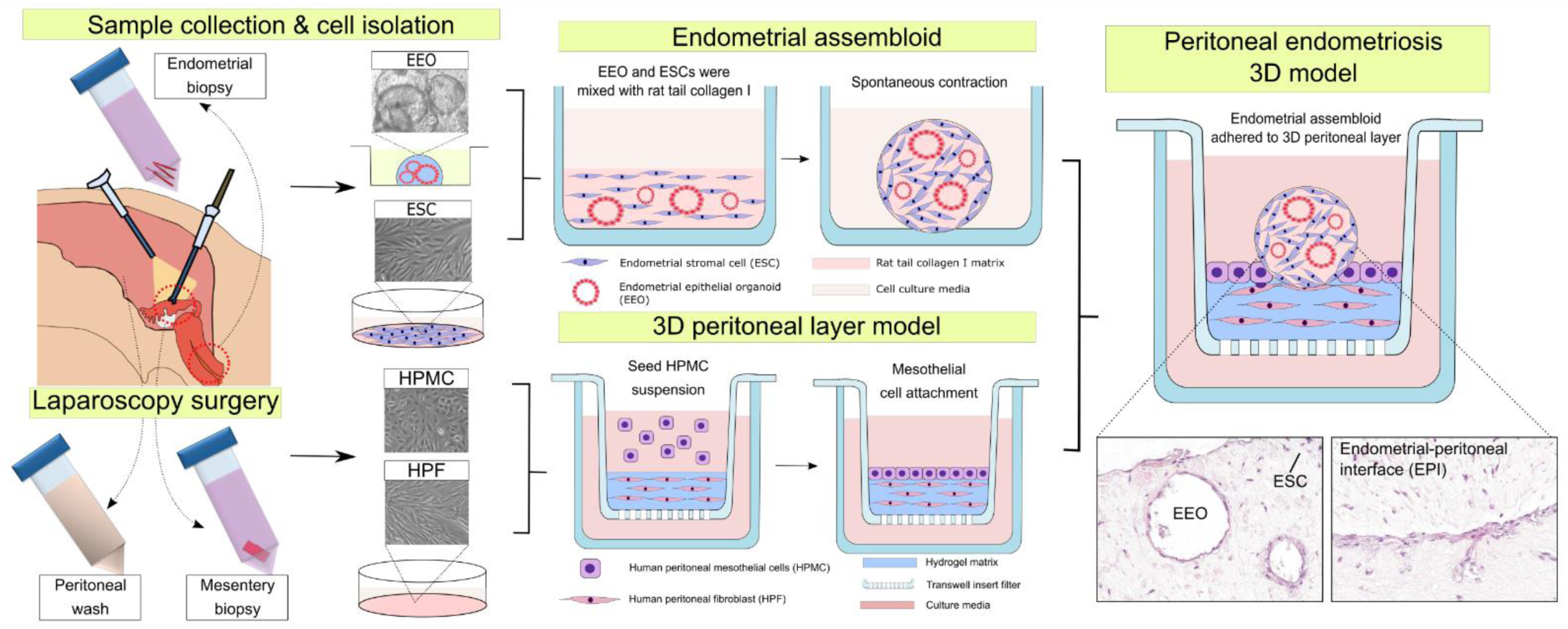

## 1. Introduction

Endometriosis is a common, incurable, chronic inflammatory condition associated with significant morbidity in at least 1 in 10 women of reproductive age, equating to over 190 million women globally [1, 2]. It typically causes debilitating chronic pelvic pain and is associated with subfertility, affecting the quality of life and wellbeing of sufferers, with an annual cost estimated at over £8.2 billion to the UK economy attributed to productivity losses and healthcare expenses [3]. The available treatments mainly comprise of hormones, which are contraceptive and associated with significant side effects, while surgical excision is risky and not curative. More effective and fertility-preserving treatments are thus urgently needed. Understanding endometriosis pathophysiology and identifying key molecular markers as targets for therapeutic interventions are essential for the development of novel treatments [4].

Traditionally, endometriosis is defined as the presence of abnormally located endometrium-like epithelial glands and stromal tissue outside of the uterus. There are three main types; (1) superficial peritoneal, (2) ovarian and (3) deep endometriosis. Superficial peritoneal endometriosis, where endometriosis lesions are located on the peritoneal surface of the pelvis, is the most common variety requiring an invasive surgical procedure (laparoscopy) for diagnosis, which is typically delayed [5]. It is also the most challenging to manage due to high recurrence rates (>70%) even after surgical excision [6]. The most accepted theory of endometriosis initiation is “Sampson’s theory”, which proposes that sloughed endometrial tissue at the time of menstruation, is transferred to the pelvic cavity via trans-tubal retrograde flow, where it attaches, invades and grows on the peritoneal surface, while retaining an endometrial-like phenotype [7, 8]. The peritoneal layer is thus postulated to play an important role in this process.

The peritoneum is composed of a simple epithelial mesothelium, basement membrane and underlying vascularised sub-mesothelial stroma containing fibroblasts and other cell types [9]. The peritoneum acts not only as a protective barrier but also interacts with macrophages within the cavity [10] and has an important role in fluid transport [11]. In response to endometriosis, peritoneal dialysis or surgical injury, the peritoneum can undergo pathophysiological changes that can lead to scarring. Both endometriosis and metastatic peritoneal disease involve the invasion of cells into the peritoneal layer, suggesting common cellular and molecular mechanisms involving mesothelial and sub-mesothelial cell behaviour [7, 12]. However, detailed analyses of the mechanisms regulating cell attachment, invasion and scar formation in the peritoneal layer are lacking in the published literature.

Endometriosis tissues retain hormone responsiveness and undergo cyclical growth, differentiation, bleeding and regeneration incurring an inflammatory response and scarring at the ectopic sites [4, 13, 14]. The associated chronic pain symptoms are particularly exacerbated at the time of menstruation and endometriosis is directly relevant to the menstruation process, which only occurs in women and upper order primates [4].

Since menstruation is irrelevant to most laboratory animals, it cannot be accurately modelled *in vivo*. Complex *in vitro* modelling using patient derived materials is thus essential to understand endometriosis aetiology and pathophysiology. The existing *in vitro* models of peritoneal endometriosis lack inclusion of both endometrial and peritoneal cell types [15, 16]; have no 3D configuration [17, 18], do not use patient-derived cells [19–21] and thus, are poor mimics of the lesions in patients.

Therefore, to overcome these deficiencies, we developed a 3D multicellular patient-derived model of early peritoneal endometriosis, containing the main endometrial and peritoneal cell types.

## 2. Materials and Methods

### 2.1 LP-9 and normal human dermal fibroblast cell culture

The human mesothelial cell strain LP-9 (AG07086) was originally isolated from the peritoneal cavity of an ovarian cancer patient and was purchased from the Coriell Institute (New Jersey, USA). LP-9 cells were cultured in mesothelial culture medium consisting of 1:1 [v/v] Medium 199 (Sigma-Aldrich, M4530)/MCDB105 (Sigma-Aldrich, 117-500). The media was supplemented with 2 mM L-glutamine (Sigma-Aldrich, G7513), 15% [v/v] foetal bovine serum (FBS, Merck, F7524), 10 ng/ml human epidermal growth factor (hEGF, Lonza, CC-4107), 0.4 µg/ml hydrocortisone (Sigma-Aldrich, H0888), and 100 µg/ml Primocin (Invivogen, ant-pm-2).

Normal human dermal fibroblasts (NHDFs, C-12300) isolated from the dermis of juvenile foreskin were purchased from Promocell (Heidelberg, Germany). NHDFs were cultured in Fibroblast Growth Medium (Promocell, C-23110) containing 1 ng/ml recombinant human basic Fibroblast Growth Factor (bFGF) and 5 μg/ml recombinant human Insulin, supplemented with 2 mM L-glutamine and 100 µg/ml Primocin. All cells were cultured on 0.1% [w/v] gelatine-coated plates at 37 °C with 5% CO_2_.

### 2.2 Ethical approval

The collection and use of human tissue was approved by the Liverpool Adults Ethics Committee (REC:19/SC/0449; approval date: 07/09/2029). All women involved provided written informed consent to the use of their tissue biopsies and relevant clinical information.

### 2.3 Biosample collection

Peritoneal wash fluid (PWF), fallopian tube mesentery, eutopic endometrium and endometriosis lesion biopsies were collected from women undergoing laparoscopic surgery for benign conditions at Liverpool Women’s Hospital. Information on participants’ demographics, samples collected, and the experiments in which their samples were utilised is presented in **Supplementary Table S1**.

### 2.4 Primary peritoneal mesothelial cell isolation and culture

Human peritoneal mesothelial cells (HPMCs) were collected from PWF following a previously described method [22] with some modifications. PWF was collected under sterile conditions during peritoneal cavity wash with saline solution and centrifuged at 300 *× g* for 10 min to obtain a cell pellet. Ficoll-Paque (Sigma, GE17-1440-02) was used to remove red blood cells from the sample. The HPMC cell pellets were frozen in Recovery Cell Culture Freezing Medium (Gibco, 12648010) and stored at −80 °C before use. HPMCs were cultured under the same conditions as LP-9 cells, with a seeding density of 1.67×10^5^ cells/cm^2^ (**Supplementary Figure S1**).

### 2.5 Primary peritoneal fibroblast cell isolation and culture

Human peritoneal fibroblasts (HPFs) were isolated from fallopian tube mesentery biopsies following a previously described method [23] with some modifications. Briefly, fallopian tube mesenteries were initially washed with sterile PBS (Gibco, 10010-015) three times, then incubated in 1x Trypsin-EDTA (Sigma, T4174) in PBS for 20 – 30 min at 37 °C in a shaking water bath to remove the mesothelial layer. This step was repeated twice before cutting the mesentery biopsies into small explants (1 – 2 mm) using scalpel blades. The explants were plated in a 6-well plate and covered with 1.5 – 2 ml of Fibroblast Growth Medium. When cells had started growing out of the explants and reached 20 – 30% confluency, the explant pieces were removed and discarded to minimise further mesothelial contamination. Medium was changed every 2 – 3 days. Upon reaching 90 – 95% confluency, HPFs were detached from the plate using 1x Trypsin-EDTA, frozen in Recovery Cell Culture Freezing Medium, and stored at −80 °C. For further expansion, HPFs were cultured under the same conditions as NHDFs with a seeding density of 1.67×10^5^ cells/cm^2^ (**Supplementary Figure S1**), except that they were cultured on non-coated plates to minimise mesothelial cell adherence.

### 2.6 Isolation of primary endometrial epithelium and stroma

Human endometrial epithelium (glands) and stroma were isolated from endometrial pipelle biopsy as previously described [24]. Endometrial tissue was cut in to small fragments (<1 mm) using a scalpel blade and further enzymatically digested with 1 mg/ml Dispase II (Gibco, 17105041), 2 mg/ml collagenase type I (Gibco, 17100017) and 80 µg/ml deoxyribonuclease (DNase) I (Merck, 11284932001) for 40 min to 1 h at 37°C in a shaking water bath. The digests were triturated every ∼15 min to facilitate tissue breakdown into free epithelial glands and single stromal cells. The endometrial digests were then filtered through a 40 µm cell strainer (Corning, CLS431750) to separate the glandular (retentate) and stromal (flow-through) components. Erythrocytes were removed from the endometrial stromal fraction using Ficoll-Paque. The intact endometrial glands and stromal cells were frozen in Recovery Cell Culture Freezing Medium and stored at −80 °C until use.

### 2.7 Endometrial epithelial organoid expansion

Endometrial epithelial organoids (EEOs) were derived from the isolated endometrial glands based on a previously published method [25] with some changes. Frozen endometrial glands were thawed at 37 °C and washed in 4 ml DMEM/Ham’s F-12 (Gibco, 21041033). Subsequently, endometrial glands were mixed with Matrigel (Corning, 536231) at 1:20 [v/v] ratio. EEOs were seeded onto a 48-well plate by placing a 20 µl-drop of Matrigel-EEO suspension, solidifying at 37 °C for 15 min, and then covering with 250 μl of organoid expansion media (**Supplementary Table S2**). The expansion media was refreshed every 2-3 days until organoids were dense. EEOs were removed from Matrigel using the Cell Recovery Solution (Corning, 354253) for endometrial assembloid generation.

### 2.8 Endometrial stromal cell culture

Endometrial stromal cells (ESCs) cells were cultured in DMEM/Ham’s F-12 media supplemented with 10% FBS and 100 µg/ml Primocin. ESCs were cultured on non-coated plates at 37 °C with 5% CO_2_, with a minimum seeding density of 3.5×10^4^cells/cm^2^.

### 2.9 Endometrial assembloid construction

Endometrial assembloids were generated by embedding EEOs and ESCs into a collagen I hydrogel (Gibco, A-10483), which was prepared according to the manufacturer’s protocol. To make a single endometrial assembloid, EEOs from one well and ESCs (4×10^4^ cells) were mixed with 100 µl collagen I at 2 mg/ml. The mixture was prepared on ice and 100 µl was deposited per well in a 96-well plate. Subsequently, the EEO-ESC-collagen mixture was set to gelate for 40 – 60 min at 37 °C. Endometrial assembloids were maintained in 200 µl of ESC culture media for 24-48 hours, at which point the assembloids had contracted and were harvested for use in the endometriosis model.

### 2.10 Peritoneal layer model construction

A 3D peritoneal layer model was generated by stepwise combination of peritoneal mesothelial cells (LP-9 or HPMCs) and fibroblasts (NHDFs or HPFs). Four different hydrogel mixtures (M1 – M4) were used in this study (**Table 1**): M1 (2 mg/ml rat tail collagen I), M2 (2 mg/ml rat tail collagen I and Matrigel matrix at 70:30 [v/v] ratio), M3 (2 mg/ml rat tail collagen I and Matrigel matrix at 50:50 [v/v] ratio and M4 (2 mg/ml rat tail collagen I supplemented with 10 µg/ml human plasma fibronectin (Merck, FC010)). Initially, fibroblasts were mixed with rat tail Collagen I-based hydrogels at 2,000 cells/µl and 100 µl of the fibroblast-hydrogel mixture was placed in the upper compartment of a 6.5 mm diameter, 3.0 µm polycarbonate membrane transwell insert (Costar, 3415). Subsequently, the transwell insert was placed in a 24-well plate and set to gelate at 37 °C for 40 – 60 min. Next, 100 µl of a cell suspension containing LP-9 or HPMCs in mesothelial culture medium (250 cells/µl) was seeded onto the upper compartment (apical) of the transwell insert, covering the fibroblast-hydrogel mixture. Fibroblast Growth Medium was added to the lower compartment (basolateral) of the well to maintain NHDF/HPF nutrients and prevent dehydration. The 3D peritoneal layer model was cultured at 37 °C with 5% CO_2_. Media was refreshed every 2 – 3 days, and the conditioned media was collected at days 1, 3, 5, 7, and 10 for analysis.

**Table 1.** Rat tail collagen I-based hydrogel mixture ratio for 3D peritoneal model construction.

| Hydrogel mixture | Collagen I<sup>1</sup> | Matrigel<sup>2</sup> | Fibronectin<sup>3</sup> |
| --- | --- | --- | --- |
| M1 | 100 | 0 | 0 |
| M2 | 70 | 30 | 0 |
| M3 | 50 | 50 | 0 |
| M4 | 99 | 0 | 1 |

<sup>1</sup>) 2 mg/ml rat tail Collagen I (Gibco, A-10483)
<sup>2</sup>) Matrigel matrix basement membrane (Corning, 536231)
<sup>3</sup>) 1 mg/ml human plasma fibronectin (Merck, FC010)

### 2.11 Peritoneal endometriosis model construction

A peritoneal endometriosis model was generated by explanting an endometrial assembloid onto the peritoneal layer model. Initially, the culture media on the apical and basolateral compartments of the peritoneal layer cultures were discarded. The endometrial assembloid was gently extracted from the 96-well plate using manual pipetting and placed onto the surface of the peritoneal layer model. The superficial endometriosis model was maintained under peritoneal layer model culture conditions.

### 2.12 Tissue biopsy and in vitro model processing for histology

Human tissue biopsies and 3D cell culture constructs were fixed in 10% neutral buffered formalin (NBF, Sigma-Aldrich, HT501128) overnight (tissue biopsies) or for 30 min (*in vitro* models) at room temperature (RT). The peritoneal layer and peritoneal endometriosis model constructs were detached from the transwell insert by cutting out the filter membrane using a scalpel blade. Constructs were set in Histogel (Fisher Scientific, 12006679) to maintain their shape during the embedding process. All tissue biopsies and the 3D models were processed to paraffin, sectioned (3 μm thickness) and placed on APES-coated glass slides for histological and immunostaining analysis.

### 2.13 Haematoxylin and eosin (H&E) staining

Sections were deparaffinised in xylene and rehydrated using an ethanol gradient. Sections were stained with Epredia™ Shandon Gill 2 Haematoxylin (Fisher Scientific, 10096648) and Epredia™ Shandon Eosin-Y (Fisher Scientific, 10188418), dehydrated in ethanol and xylene, and mounted in Epredia™ Consul-Mount™ medium (Fisher Scientific, 9990441).

### 2.14 Immunohistochemistry (IHC) staining

IHC was conducted as described previously [26]. In short, deparaffinised sections underwent antigen retrieval and were incubated with primary antibodies (**Supplementary Table S3**) overnight at 4 °C. Sections were then washed thoroughly in tris-buffered saline (TBS) before applying horse radish peroxidase-conjugated ImmPRESS anti-mouse IgG (Vector Laboratories, MP-7402) or anti-rabbit IgG (Vector Laboratories, MP-7401) secondary antibodies for 30 min at RT. Antibody detection was visualised by incubating the sections for 10 min at RT with 3, 3’-diaminobenzidine (DAB) (ImmPACT Vector Laboratories, SK-4105). Samples were counterstained using haematoxylin, dehydrated and mounted as described above.

### 2.15 Immunofluorescence (IF) staining

LP-9, NHDFs, HPMCs and HPFs were cultured in 8-well Nunc™ Lab-Tek™ II Chamber Slides™ (Thermo Scientific, 154534) at a seeding density of 15,000 - 20,000 cells/well, at 37 °C until 80–90% confluency. Cells were fixed in –20 °C absolute ethanol for 10 min followed by two gentle PBS washes. Cells were permeabilised with 0.25% [v/v] Triton X-100 (Thermo Scientific, 85111) in PBS for 10 min at RT and blocked with 2% [w/v] bovine serum albumin (BSA, Sigma-Aldrich, A3803) for 1 h at RT. Cells were incubated with primary antibodies overnight at 4 °C (**Supplementary Table S3**), followed by three 15 min PBS washes and incubation with secondary antibodies AlexaFluor™ 488 goat-anti-mouse IgG1 (Invitrogen, A21121) and AlexaFluor™ 568 goat-anti-rabbit (Invitrogen, A11011) at 1:1,000 dilution, and DAPI (Sigma, D9542) in a 1:500 dilution, for 1 h at RT. The slides were mounted using Fluoromount™ media (Invitrogen, 00-4958-02). For IF staining of the 3D peritoneal layer model, sections were prepared and treated with primary antibodies (**Supplementary Table S3**) and AlexaFluor™-labelled secondary antibodies with DAPI at concentrations mentioned above. An Autofluorescence Quenching Kit (Vector Laboratories, SP-8400-15) was applied for 5 – 10 min at RT prior to imaging.

### 2.16 Lactate dehydrogenase assay

To assess cell viability, lactate dehydrogenase (LDH) release was quantified in conditioned media of the mixed basolateral and apical compartments of the transwell insert using an LDH cytotoxicity assay kit (Roche, 11644793001 v12). Samples were assayed in duplicate following the manufacturer’s protocols. Optical density was measured at 490 nm using a FLUOstar Omega microplate reader (BMG Labtech) and fresh culture media as control.

### 2.17 Enzyme-linked immunosorbent assay (ELISA)

Human tissue plasminogen activator (tPA) was measured in the conditioned medium obtained from the apical compartment of the 3D cell culture system using a human tPA DuoSet ELISA kit (Biotechne, DY7449-05), following the manufacturer’s protocol. Standard curves (**Supplementary Figure S2**) were generated from 2-fold serial dilutions of tPA standard solution. Samples were assayed in duplicate by measuring optical density at 450 nm using a FLUOstar Omega microplate reader as previously described [27].

### 2.18 Imaging and data analysis

Stained sections were digitalised using an Aperio CS2 slide scanner (Leica Biosystems, 23CS100CE) and images were visualised using Aperio ImageScope Version 12.4.6.5003. Cell shape of growing cultures was recorded using bright field microscopy (Leica DM IL), while IF-stained cells were visualised using epifluorescence microscopy (Leica DM2500). Images were processed with Fiji [27]. Quantification of cell marker expression was performed by manually counting positive cells in 3 – 5 regions of interest per well using the Fiji Cell Counter plugin and subsequently calculating the mean percentage ± standard deviation (SD). Axial view images of peritoneal models were acquired using a dissection microscope (Leica MZ16 F). The average endometrial assembloid diameter ± SD was measured using Fiji. Standard curve of tPA ELISA was generated in GraphPad Prism Version 9.0 using a four-parameter logistic regression model. Data were visualised as graphs using OriginPro 2024b.

## 3. Results

### 3.1 Isolated primary peritoneal mesothelial cells and fibroblasts demonstrate cell type-specific characteristics

Peritoneal wash-derived human peritoneal mesothelial cells (HPMCs) and fallopian tube mesentery-derived human peritoneal fibroblasts (HPFs) were initially characterised by phase-contrast microscopy to distinguish their morphology. Morphological characteristics of HPMCs and HPFs were compared to the normal human mesothelial cell line LP-9, and primary normal human dermal fibroblasts (NHDFs), respectively, in a two-dimensional (2D) cell culture system. At passage 2 (P2), HPMCs had a cobblestone appearance comparable to that of LP-9 cells at P7, demonstrating their expected mesothelial morphology. HPFs at P1 exhibited a spindle-shape appearance comparable to NHDFs at P11 (**Figure 1A**). To confirm peritoneal mesothelial and fibroblast cell phenotypes, dual immunofluorescence staining of cytoskeletal markers cytokeratin (CK) and vimentin was performed in cells (**Figure 1B**) [28]. Our findings indicated that CK and vimentin were localised in the cytoplasm of the peritoneal mesothelial cells, while the NHDFs and HPFs expressed vimentin and lacked CK. Furthermore, scoring for the ratio of CK and vimentin expression in six biological HPMC samples showed that the HPMCs at P2 had a very high percentage of CK^+^/vimentin^+^ cells (mean 98.20 ± 1.05 SD). Meanwhile, isolated HPFs at P2 from three biological samples exhibited CK^−^/vimentin^+^ expression of 81.36 ± 5.63% cells, indicating their fibroblast characteristics. HPFs also contained 18.64 ± 5.63% CK^+^/vimentin^+^ cells, suggesting a minor contamination with mesothelial cells (**Figure 1B**).

**Figure 1.**
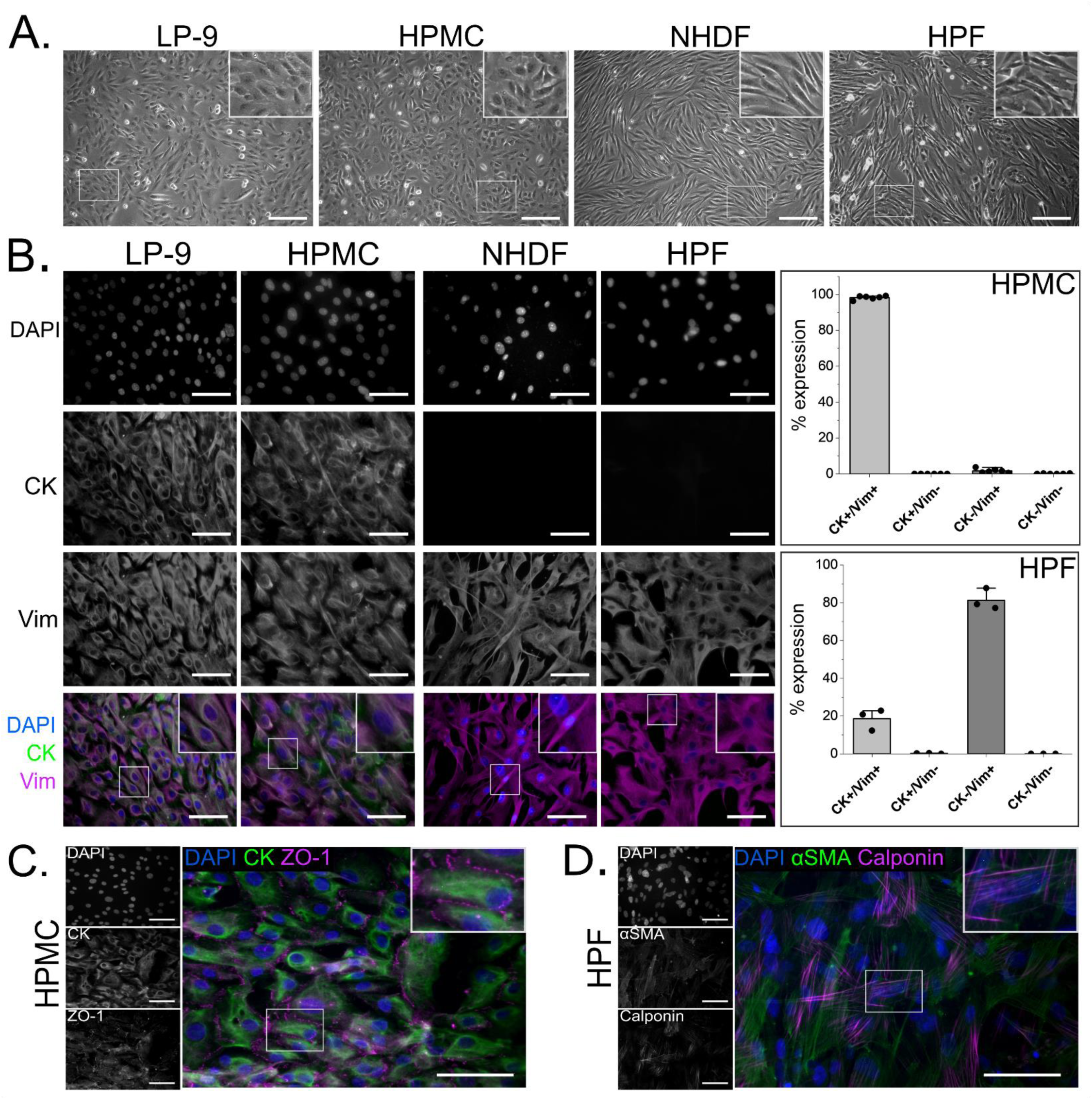
Characterisation of primary human peritoneal mesothelial cells (HPMCs) and human peritoneal fibroblasts (HPFs) compared to the LP-9 mesothelial cell line and normal human dermal fibroblasts (NHDFs). (A) Representative phase contrast micrographs of LP-9, HPMCs, NHDFs and HPFs. Mesothelial and fibroblast cells exhibit cobblestone and spindle-like morphologies, respectively. (B) Antibodies against the cytoskeletal markers cytokeratin (CK) and vimentin (Vim) were localised in LP-9 cells and HPMCs, while only vimentin but not CK was expressed in NHDFs and HPFs. (C) Expression of tight junctional marker Zonula Occludens-1 (ZO-1) was detected at HPMC junctions. (D) Isolated HPFs expressed various levels of α-smooth muscle actin (αSMA) and calponin stress fibres. Scale bars: A, 200 µm; B-D, 50 µm.

Epithelial characteristics of isolated HPMCs were confirmed by uniform expression of the tight junction marker Zonula Occludens-1 (ZO-1) in the cell membrane (**Figure 1C**) [28]. Further characterisation of HPFs revealed expression of α-smooth muscle actin (αSMA) and calponin as stress fibres in the cytoplasm (**Figure 1D**), suggesting that the isolated HPFs in 2D culture were in various physiological states [29].

### 3.2 Generating a composite 3D hydrogel construct using patient-derived peritoneal cells in vitro

An *in vitro* peritoneal layer model was generated by incorporating peritoneal mesothelial cells and fibroblasts in a 3D co-culture system (**Figure *2*A**). Initial experiments were performed using commercially available cells (LP-9 and NHDFs) before trialling patient-derived cells (HPMCs and HPFs). A hydrogel-fibroblast mixture (NHDFs or HPFs) mimicking the sub-mesothelial layer was placed in the upper compartment of a transwell insert. This design allowed sufficient access to nutrients and supported the matrix to remain flat [30]. Mesothelial cells (LP-9 or HPMCs) were then seeded onto the surface of the hydrogel-cell matrix to mimic the mesothelial layer. In our approach, we used HPMCs and HPFs from the same patients to ensure compatibility of the cells within the constructs (**Supplementary Table 1**). We cultured the composite hydrogel constructs for up to 10 days and analysed the histological structure of the constructs on days 3 and 10. Axial observation of composite hydrogel constructs revealed the formation of flat-shaped disks *in vitro* (**Figure 2B**). Histological analysis revealed that a single cell layer covered the surface of the hydrogel-cell matrix, suggesting the formation of a mesothelial layer by day three. The hydrogel matrix contained embedded cells mimicking the sub-mesothelial stroma (**Figure 2B**).

**Figure 2.**
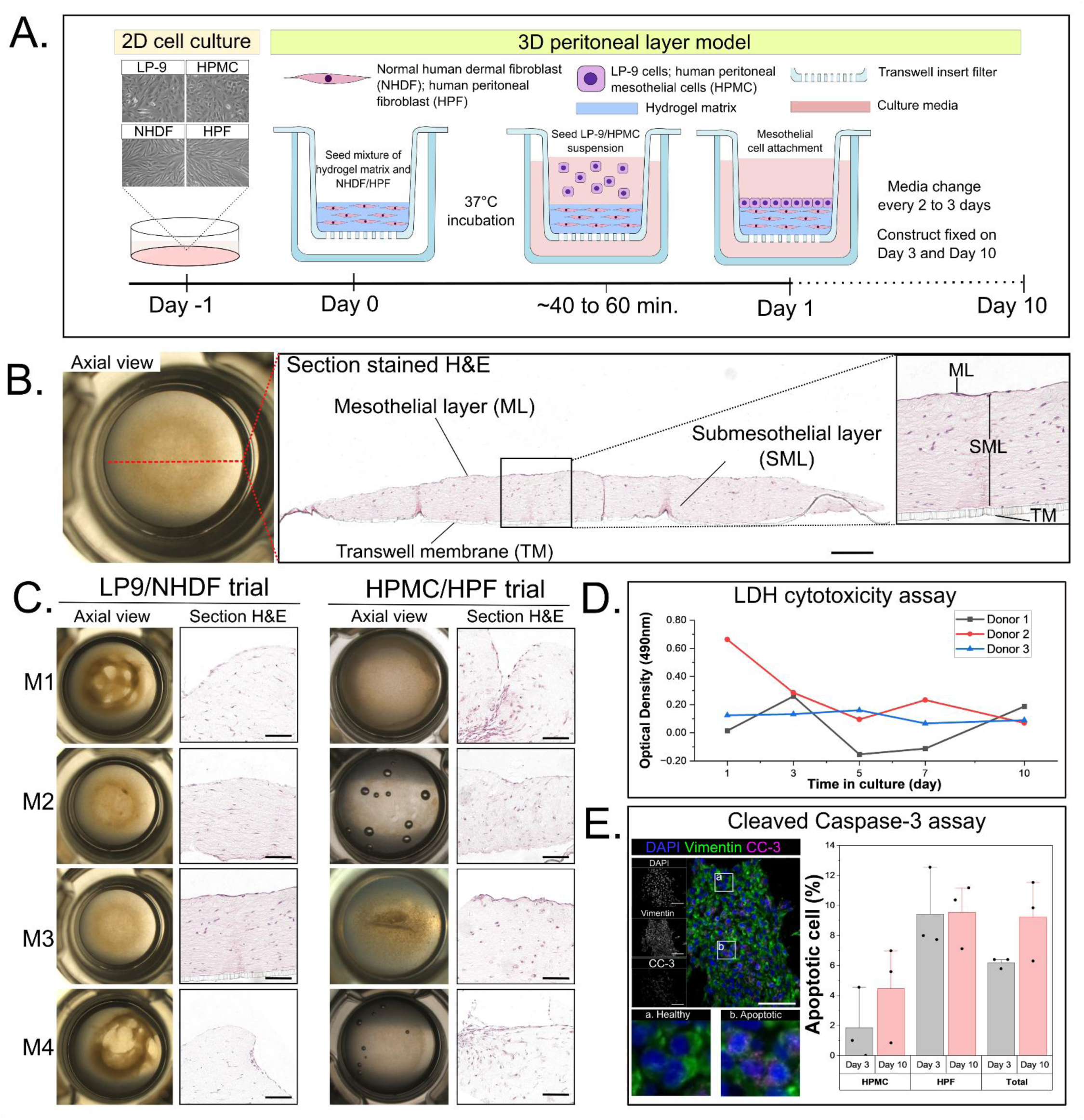
Establishing composite 3D hydrogel constructs using peritoneal mesothelial cells and fibroblasts. (A) Schematic illustration of model construction and culture timeline. (B) Representative axial view and H&E-stained section of a hydrogel construct showing the formation of a mesothelial monolayer (ML) and sub-mesothelial layer (SML) on a transwell membrane (TM). (C) Representative images of hydrogel matrices using M1 (collagen I), M2 (70:30 collagen I:Matrigel ratio), M3 (50:50 collagen I:Matrigel ratio) and M4 (collagen I + human fibronectin). Construct generated with matrix combination M3 demonstrated minimal contraction in LP-9/NHDF and HPMC/HPF trials. (D) Lactate dehydrogenase (LDH) cytotoxicity assay in M3 composite hydrogel constructs containing HPMC/HPF (*n* = 3 donors) over a 10-day culture period. (E) Dual immunofluorescence staining of cleaved caspase-3 (CC-3) and vimentin to detect apoptotic HPMCs/HPFs in M3 constructs on day 3 and day 10 of culture (*n* = 3). Scale bars: B, 300 µm; C, 100 µm; E, 50 µm.

To determine the optimal composition of the hydrogel scaffold, four different combinations were trialled: M1 (collagen I), M2 (70:30 collagen I:Matrigel), M3 (50:50 collagen I:Matrigel) and M4 (collagen I + human fibronectin). Composite hydrogel cultures containing LP-9/NHDF or HMPC/HPF combinations demonstrated differing levels of contraction on day three post-formation (**Figure 2C**). Histological analysis of transverse sections indicated that different matrix compositions affected the structural appearance of the hydrogel constructs; the upper contour of the constructs varied between matrix combinations. However, matrix combination M3 provided the best support in establishing an *in vitro* model of the peritoneal layer with minimal contraction using both commercial cells (LP-9 and NHDFs) and patient-derived cells (HPMCs and HPFs). Therefore, the combination M3 consisting of equal parts collagen I and Matrigel was taken forward.

To assess cell viability within the composite hydrogel over the experimental period, conditioned media from the basolateral and apical compartments of transwell inserts were subjected to LDH cytotoxicity analysis. LDH release remained relatively static across 10 days of culture in constructs composed of cells from three donors (**Figure *2*D**). A high level of LDH release was observed on day one of culture in donor 2, likely due to cell damage induced following model generation. However, LDH release was comparable across donors by day three. The number of apoptotic cells within the constructs was quantified by cleaved caspase-3 expression (**Figure 2E**) HPMCs and HPFs were highly viable in the hydrogel constructs, with overall 5.92 ± 0.42%, and 9.22 ± 2.67% of apoptotic cells found at 3 and 10 days of culture, respectively. These findings suggested that the composite 3D hydrogel construct is suitable for long-term culture of patient-derived peritoneal cells.

### 3.3 The composite 3D hydrogel construct is a simple mimic of the peritoneal layer in architecture and function

The microstructure of the composite 3D hydrogel construct was assessed using histological staining on longitudinal paraffin-fixed sections. The construct was compared to patient-derived parietal peritoneum biopsies from the uterovesical fold to determine how closely it resembled the histological and phenotypical features of the peritoneum. Histological analysis of constructs generated using LP-9/NHDFs or HPMCs/HPFs revealed a cell monolayer on their surface and a sparser layer of cells underneath, thus recapitulating the mesothelial and sub-mesothelial layers of the parietal peritoneum (**Figure 3A**). Cytokeratin and podoplanin were strongly expressed in the surface layer of the construct similar to the mesothelial layer of the parietal peritoneum, confirming their mesothelial origin (**Figure 3A**) [31]. As in the parietal peritoneum, we also detected collagen IV expression in a thin layer beneath the mesothelium in the constructs, indicating the spontaneous formation of a basal lamina by LP-9 and HPMC monolayers (**Figure 3A**). Cells within the bipartite matrix expressed fibroblast specific protein-1 (FSP-1) and α-smooth muscle actin (αSMA), which was comparable to the sub-mesothelial layer of the parietal peritoneum (**Figure 3B**). However, some mesothelial cells in the parietal peritoneum also expressed FSP-1, suggesting variability of fibroblast protein expression levels in mesothelial cells *in vivo*. LP-9 cells also expressed FSP-1 and αSMA, suggesting their upregulation under these culture conditions. Interestingly, LP-9 cells cultured in 2D under different media compositions led to variation in FSP-1 and αSMA expression (**Supplementary Figure S3**), suggesting that FSP-1 and αSMA expression in LP-9 cells are influenced by the culture environment. Together, these results indicate that LP-9 mesothelial cells may not be suitable to model a peritoneal layer *in vitro*, and patient-derived HPMCs and HPFs were used in further analysis of composite 3D hydrogel constructs as simple peritoneal layer models.

**Figure 3.**
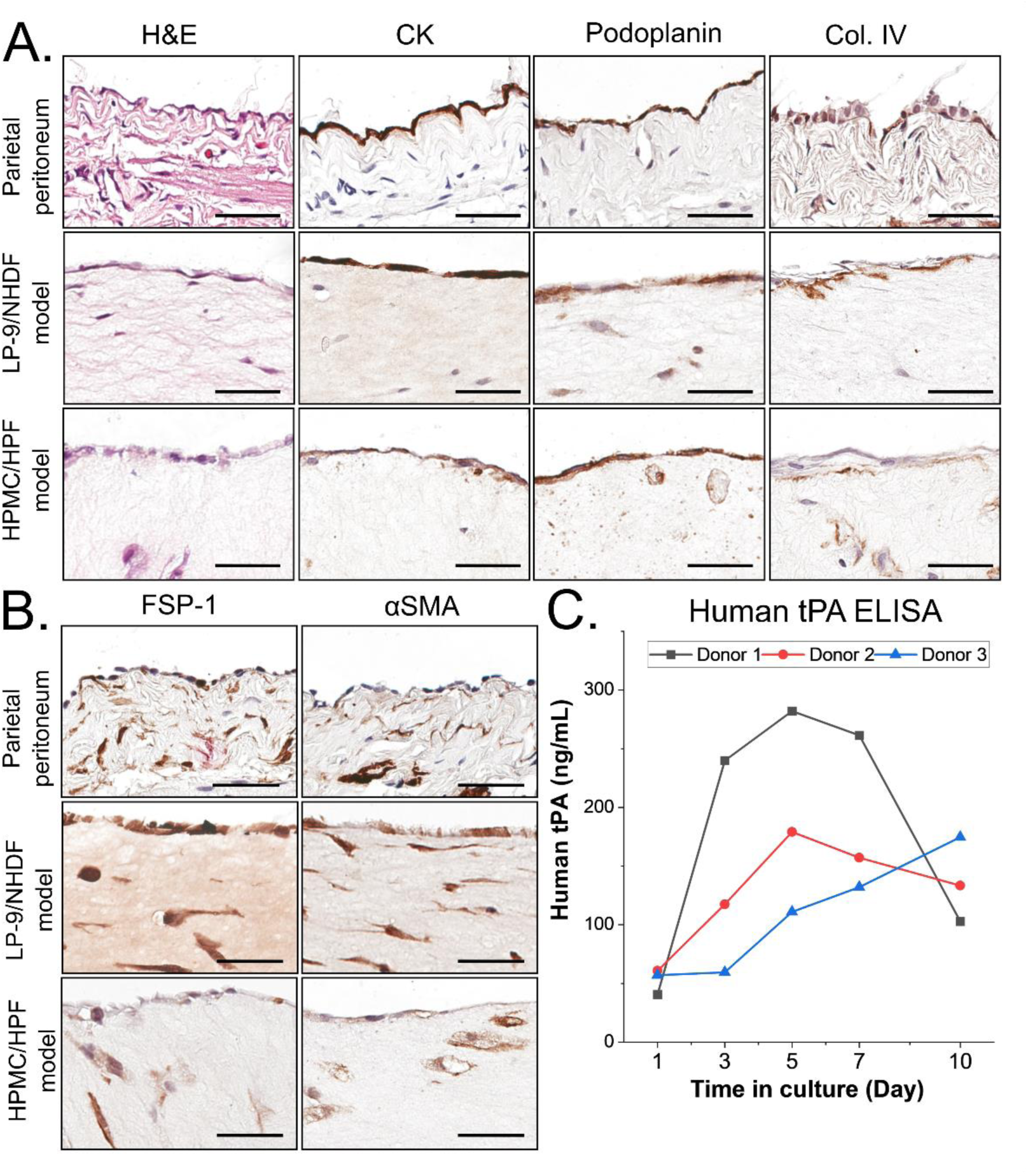
Histological and functional analysis of peritoneal layer models. (A-B) Histological staining of transverse sections through parietal peritoneum and composite 3D hydrogel constructs composed of LP-9/NHDFs and HPMCs/HPFs. Immunohistochemical staining used (A) antibodies against the cytoskeletal markers cytokeratin (CK) and podoplanin as mesothelial markers; collagen type IV (Col IV) as basal lamina marker; and (B) fibroblast specific protein-1 (FSP-1) and α-smooth muscle actin (αSMA) as sub-mesothelial markers. (C) Human tissue plasminogen activator enzyme-linked immunosorbent assay (tPA ELISA) to determine the functionality of the mesothelial cells in models assembled with matched cells from three different donors over a 10-day culture period. Scale bars: A-B, 100 µm.

To determine the functionality of the composite 3D hydrogel construct as a peritoneal layer model, we assessed the level of human tissue plasminogen activator (tPA) [32, 33] in conditioned media from the apical compartments of transwell inserts (**Figure 3C**). Human tPA was detected from day 1 over the 10-day culture period using matched cells from three different donors, with an average peak on day 5 (190.68 ± 81.13 ng/ml), in line with reported tPA levels in cultured mesothelial cells [32]. These results indicate normal physiological behaviour of HPMCs in the peritoneal layer model.

### 3.4 Endometrial assembloids show similar characteristics to the eutopic endometrium

To constitute the endometrial component of superficial endometriotic lesions, endometrial assembloids were generated from patient-derived cells. Endometrial epithelial organoids (EEOs) and endometrial stromal cells (ESCs) were combined in a collagen I hydrogel that spontaneously contracted to form spherical assembloids with an average diameter of 1.15 ± 0.07 mm (**Figure 4A, B**). Histological analysis of assembloids confirmed retention of endometrial architecture and phenotype (**Figure 4C**); EEOs expressed cytokeratin and retained a luminal space within the assembloids. ESCs expressed CD10 and were distributed throughout the assembloids and formed cell-cell interactions with EEOs (**Figure 4D**).

**Figure 4.**
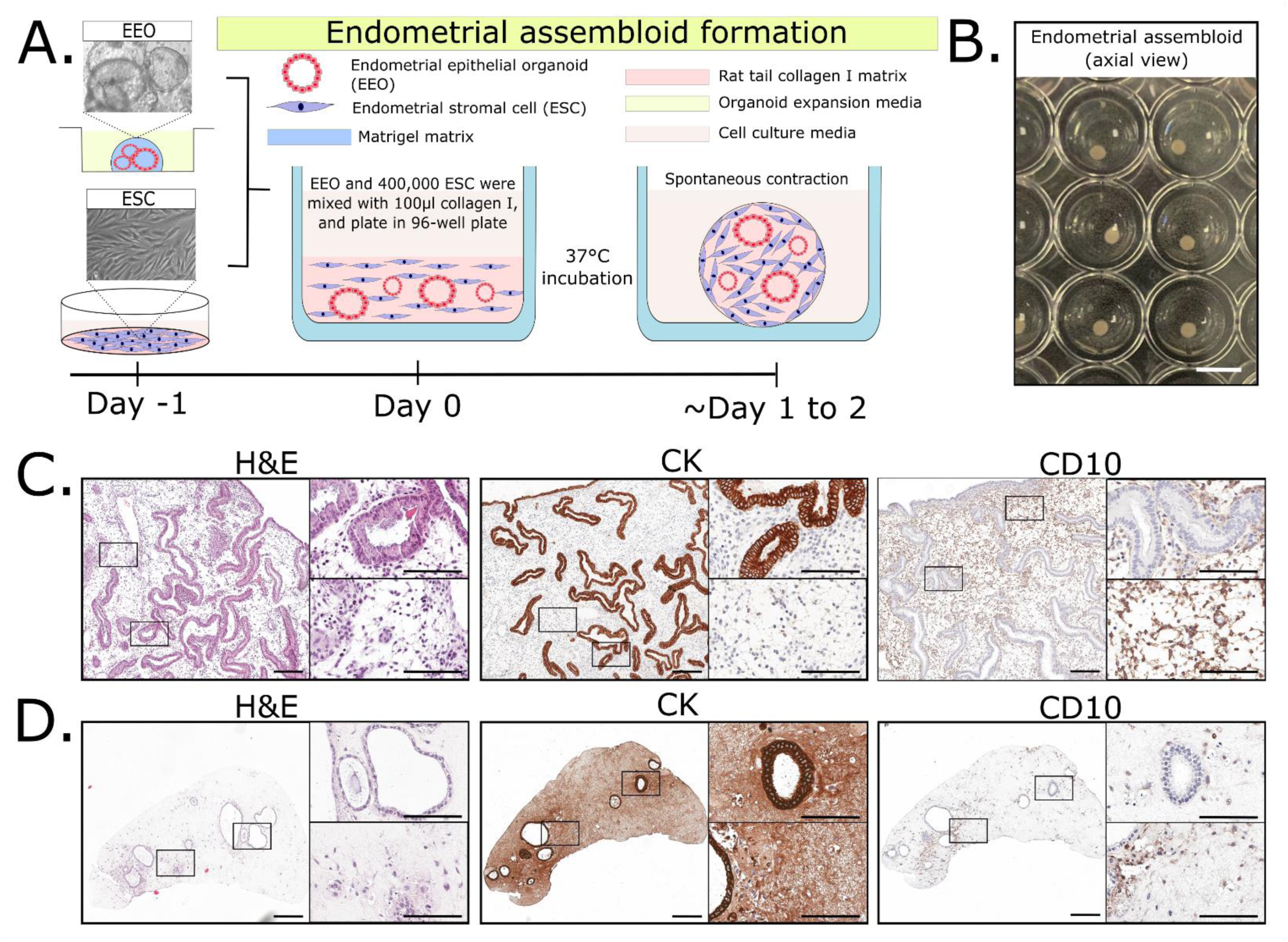
Assembly and characterisation of endometrial assembloids from endometrial epithelial organoids and endometrial stroma. (A) Schematic illustration of endometrial assembloid formation. (B) Representative axial view of contracted endometrial assembloids in a 96-well plate. (C-D) Histological staining of the cytoskeletal markers cytokeratin (CK) and CD10 on transverse sections of the eutopic endometrium (C) and endometrial assembloids (D). Scale bars: B, 3.2 mm; C – D, 200 µm (zoom-out images), 100 µm (zoom-in images).

### 3.5 Endometrial assembloids can be implanted onto the peritoneal layer model to mimic early superficial endometriosis formation

To create an *in vitro* model of superficial endometriosis, endometrial assembloids were transplanted onto peritoneal layer models (**Figure 5A**). The combined models were analysed on days 3 and 10 of co-culture. Overall model architecture revealed the fusion of endometrial and peritoneal elements (**Figure 5B**); endometrial assembloids were adhered to the apical surface of peritoneal layer models, creating an endometrial-peritoneal interface (EPI). EEOs within endometrial assembloids displayed gland-like structures, whilst ESCs retained expression of CD10 throughout the culture period, thus mirroring endometriotic lesion characteristics (**Figure 5C**). The EPI was composed of cells expressing cytokeratin and CD10, suggesting an interaction between HPMCs and ESCs on day 3 of co-culture. However, the mesothelial cell boundary was disrupted by day 10, with a higher abundance of CD10^+^ ESCs at the EPI (**Figure 5C**). These findings indicate the migration of ESCs from the endometrial to the peritoneal element of the model, thus recapitulating endometriotic lesion establishment *in vitro*.

**Figure 5.**
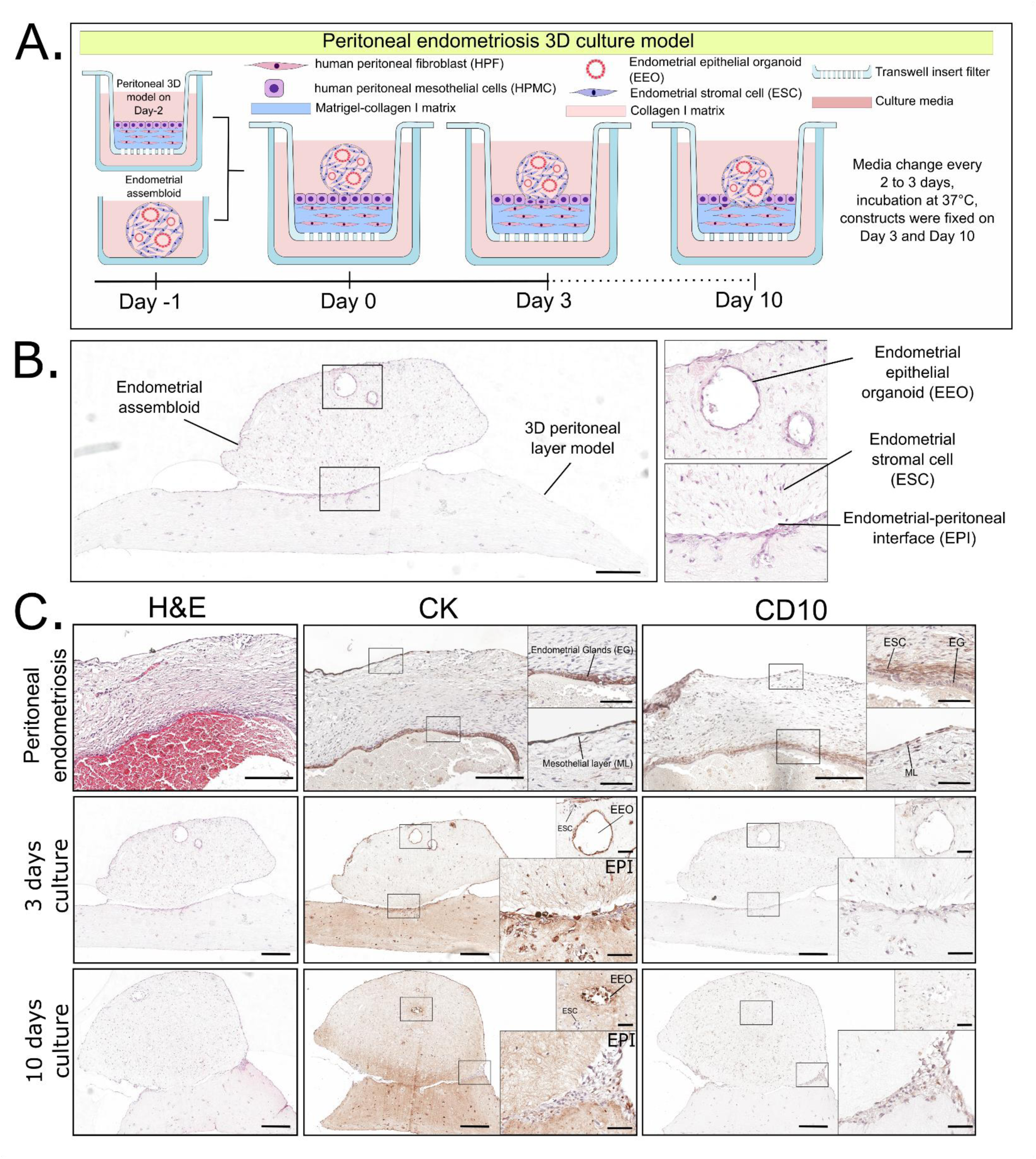
Characterisation of the superficial endometriosis model. (A) Schematic illustration of superficial endometriosis model assembly, comprising an endometrial assembloid and peritoneal layer model. (B) H&E stained transverse section of a superficial endometriosis model highlighting endometrial and peritoneal components. (C) Histological staining of superficial endometriosis lesions and in vitro model after 3 and 10 days of culture. Scale bars: B, 300 µm; C, 150 µm (zoom-out images), 50 µm (zoom-in images).

## 4. Discussion

In this study, we have demonstrated that patient-derived primary cells can be utilised to establish a 3D cell culture system modelling peritoneal endometriosis. We have observed cell-to-cell and cell-to-matrix interactions during the process of establishing a multicellular complex model, where endometrial assembloids attach to and interact with the peritoneal layer model. Importantly, our cell isolation technique is pragmatic as all needed biosamples are easily harvested from a single patient undergoing routine laparoscopic surgery without the need for any additional or high-risk invasive procedures that compromise patient safety. Peritoneal wash fluid, excised peritoneum and endometrial biopsies are common byproducts of routine surgeries undertaken in women with endometriosis, and provided the materials required to generate a multicomponent model of peritoneal endometriosis. Therefore, this study supports future development towards more ethically sound clinical applications [34] and personalised disease modelling.

The peritoneum plays an important role as the first contact point for endometrial tissue fragments in endometriosis lesion initiation based on the retrograde menstruation theory. Therefore, the peritoneal model is of great value in further exploring the cellular and molecular mechanisms involved in early endometriosis formation, which can be utilised to develop preventative strategies in the future.

With regards to the advancement of 3D cell culture techniques, a recent study has generated an *in vitro* endometriosis model by combining spheres of endometrial epithelial cells with a mesothelial monolayer. However, commercial cell lines were used, which often fail to recapitulate the phenotypic and functional characteristics of primary cells [16]. In this study, we generated a co-culture system with a peritoneal element composed of both mesothelial and submesothelial layers from patient-derived cells. Moreover, our endometrial model comprised of EEOs surrounded by ESCs, closely resembling the eutopic endometrium and endometrial fragments in menstrual effluent [35]. Our novel model recapitulates both peritoneal and endometrial architecture and is replicable and expandable.

The challenge in developing a patient-derived *in vitro* peritoneal layer model was to maintain a flat-shaped structure to be readily implanted with multicellular mimics of endometrial fragments. Our model provided means to overcome the above and allowed the process of endometrial and peritoneal interaction to be fostered for a 10-day period. Since collagen I is the most abundant extracellular matrix protein in the sub-mesothelial layer [36, 37], the utilisation of collagen I-based hydrogels as extracellular matrix mimetics could mimic the native cellular environment and support fibroblast viability *in vitro*. However, the fibroblast-collagen assortment often resulted in hydrogel contraction, which disrupts the mesothelial component [38]. Previous study suggested that the fibroblast-collagen mixture contraction could be minimised vertically using alginate cross-link and horizontally by polydopamine (PDA) adhesion [39]. Our study found that a bipartite collagen I-Matrigel matrix also minimised gel contraction in the peritoneal layer model. The use of Matrigel increases hydrogel stiffness due to the formation of larger pores that increase rigidity [40]. The contribution of Matrigel to overall hydrogel composition was limited to a 50:50 [v/v] ratio, since Matrigel is more commonly used to model cancer pathogenesis. Therefore, increasing Matrigel ratio may minimise the resemblance of our 3D peritoneal layer model to imitate the healthy peritoneal extracellular matrix. Further study is required to analyse the rheological properties of the peritoneal layer model with comparison to healthy peritoneal tissue.

The methods we have developed in biosample procurement and 3D culture systems allow further application beyond superficial endometriosis modelling; other endometriosis subtypes, such as deep endometriosis and ovarian endometriosis, may be modelled in the future. As the peritoneum is instrumental in the development of other pathologies, such as cancer metastasis, the peritoneal layer model can be utilised for diverse disease modelling pursuits. Ovarian cancer metastasis has been previously modelled by incorporating patient-derived omental adipocytes and various cell lines such as MeT5A mesothelial cells, MRC-5 fibroblasts, EA.hy926 endothelial cells and THP-1 macrophages [30]. Our patient-derived model can be expanded with the addition of immune cells and endothelial cells to study conditions such as metastasis. Further work that will benefit from the prototype we present include studies exploring trans-mesothelial water permeability in peritoneal dialysis [41] and those examining inflammation [42]. Besides, our model secreted tPA which activates plasminogen in the serosal cavity and plays important roles in peritoneal adhesion [11]. This function has been previously modelled to explore wound healing and fibrotic processes using MeT5A mesothelial cells without tPA assessment [43], suggesting the utility of our model to improve this study in the future. Therefore, our peritoneal layer model is a promising platform to be modified into a more complex mimic of *in vivo* lesions, using patient-derived primary cells and, would be applicable to other diseases apart from endometriosis.

Our superficial endometriosis 3D model allowed us to capture the EPI and characterise its composition. This creates an opportunity for future studies to explore the mechanisms of early endometriotic lesion establishment. Since tissue analysis technology continues to rapidly improved, the molecular mechanisms involved at the EPI can further be studied through spatial transcriptomics, as has previously been performed in actual endometriosis lesions and the eutopic endometrium [44, 45]. Moreover, our study demonstrated the utility of conditioned media to assess cell viability and mesothelial functioning in the peritoneal layer model. Therefore, this platform enables further study to assess cytokines, chemokines and growth factors involved in early endometriosis formation using the conditioned media, as previously assessed in endometriosis patient samples [46]. However, such analyses would be improved by incorporating immune cells into the superficial endometriosis model, since endometriosis is a complex disease involving the peritoneal immune microenvironment [47].

Regarding limitations, this study was conducted using peritoneal fibroblasts with some degree of mesothelial cell contamination, which may interfere with the cellular composition of the sub-mesothelial layer. The purity of the peritoneal fibroblast isolates could be improved by cell sorting, however, markers to differentiate mesothelial cells from fibroblasts is problematic since mesothelial cells in culture can undergo reversible spontaneous mesothelial to mesenchymal transition, especially following long-term culture [28, 48, 49]. We have not performed functional assessment of our early endometriosis model in this study, which requires further exploration in future studies. Specifically, to assess the impact of ovarian sex hormones exposure to the model as previously described [16]. Deep characterisation of the model in comparison to early endometriosis lesions using omics technologies will also prove that our peritoneal endometriosis model recapitulates *in vivo* lesions in composition and physiological functioning.

## 5. Conclusions

Peritoneal superficial endometriosis causes debilitating chronic pelvic pain and other symptoms that negatively affect the wellbeing and productivity of millions of women of reproductive age worldwide. There is no curative treatment, and diagnosis requires invasive surgical interventions. Understanding endometriosis pathophysiology and identifying diagnostic and therapeutic targets are hampered by the challenges in examining disease initiation and progression. Therefore, a physiomimetic patient-derived *in vitro* model that accurately mimics the lesions of peritoneal endometriosis is urgently needed. Such a model will overcome the bottleneck that currently exist in the endometriosis discovery pathway. Herein, we present an *in vitro* model containing the main cell types of both the peritoneal and endometrial components of endometriotic lesions, which provides a robust platform for further refinement for the benefit of endometriosis research.

## CRediT authorship contribution statement

**Muhammad Dimas Reza Rahmana**: Methodology, validation, formal analysis, investigation, resources, writing – original draft, writing – review & editing, visualization, project administration. **Christopher J. Hill**: Conceptualization, supervision, methodology, validation, formal analysis, investigation, resources, writing – original draft, writing – review &editing, project administration. **Bettina Wilm**: Conceptualization, supervision, methodology, resources, writing – original draft, writing – review & editing. **Dharani K. Hapangama**: Conceptualization, supervision, methodology, resources, writing – original draft, writing – review & editing.

## Acknowledgements

The authors gratefully acknowledge those patients who consented to the collection and use of their biosamples for research. We thank Dr Nicola Tempest and Dr Alison Maclean for biosample collection, technical team members Dr Alan Carter, Dr Joni Roachdown and Dr Rafah Al-Nafakh for biosample processing. MDRR was supported by the Centre for Higher Education Funding and Assessment, Ministry of Higher Education, Science, and Technology of Republic Indonesia, co-partner with Indonesian Endowment Fund for Education (LPDP). DKH and CJH were supported by a Wellbeing of Women’s project grant (RG2137) and an Impact Acceleration Account award from EPSRC and MRC.

## Data availability

Data will be made available on request.

