## Supplementary for "Compound design of a patient-derived 3D cell culture system modelling early peritoneal endometriosis"

*Joint senior authors

### Supplementary Tables and Figures

**Table S1.** Participant demographic

| No. | Menstrual phase | Age | BMI | Parity | Reason for surgery | Sample collected | | | | | Experiments |
|---|---|---|---|---|---|---|---|---|---|---|---|
| | | | | | | PWF | FTM | EB | UVF | ECT | |
| 1 | HT | 23 | 49.9 | 0 | Diagnostic laparoscopy | ✓ | | | | | IF |
| 2 | Secretory | 28 | 33.2 | 0 | Diagnostic laparoscopy | ✓ | | | | | IF |
| 3 | Proliferative | 34 | 25.2 | 2 | Laparoscopic sterilisation | ✓ | | | | | IF |
| 4 | HT | 28 | 27.3 | 1 | Diagnostic laparoscopy +/- excision of endometriosis | ✓ | | | | | IF |
| 5 | Mid-phase | 21 | 24 | 1 | Ovarian cystectomy | ✓ | | | | | IF |
| 6 | Secretory | 42 | 33.1 | 5 | Diagnostic laparoscopy | ✓ | | | ✓ | | IF, IHC |
| 7 | Proliferative | 37 | 29 | 3 | Other | ✓ | ✓ | | | | IF,<br>3D peritoneal model – donor 3 LDH assay and ELISA |
| 8 | Menopause | 55 | 31.5 | 2 | Other | | ✓ | | | | IF |
| 9 | Secretory | 38 | 20.5 | N/A | N/A | ✓ | ✓ | | | | 3D peritoneal model - apoptosis assay |
| 10 | Menstrual period | 33 | 32.8 | 2 | Salpingectomy + hysterectomy | ✓ | ✓ | | | | IF,<br>3D peritoneal model - apoptosis assay |
| 11 | Proliferative | 30 | 25.1 | 4 | Tubal ligation | ✓ | ✓ | | | | 3D peritoneal model – donor 2 LDH assay and ELISA, apoptosis assay<br>3D endometriosis model - H&E, IHC |
| 12 | Secretory | 23 | 30 | 0 | Diagnostic laparoscopy | | | ✓ | | | 3D endometriosis model - H&E, IHC |
| 13 | HT | 39 | N/A | 1 | N/A | ✓ | | | | | 3D peritoneal model – donor 1* LDH assay and ELISA, apoptosis assay<br>3D endometriosis model - H&E, IHC |
| 14 | Secretory | 43 | 24.5 | 0 | Menorrhagia | | | ✓ | | | IHC |
| 15 | Proliferative | 27 | N/A | 2 | N/A | | | | | ✓ | IHC |
| 16 | Secretory | 26 | 41 | 0 | Diagnostic laparoscopy | | | ✓ | | | Endometrial spheroid model |

HT (on hormone treatment); N/A (data not available) PWF (peritoneal wash fluid); FTM (fallopian tube mesentery); EB (endometrial biopsy); UVF (uterovesical fold biopsy); ECT (ectopic lesion biopsy); IF (immunofluorescence staining); IHC (immunohistochemistry staining); LDH (lactate dehydrogenase); ELISA (enzyme-linked immunosorbent assay); H&E (Haematoxylin and Eosin). *Donor 1 peritoneal layer model used peritoneal fibroblast from patient no. 11.

**Table S2.** Organoid expansion media used for endometrial epithelial organoid (EEO) expansion

| Component | Manufacturer | Catalogue reference | Final concentration |
|---|---|---|---|
| DMEM/F12 Phenol-free | Gibco | 21041025 | Not applicable |
| N2 supplement | Gibco | 17502048 | 1X |
| B27 supplement | Gibco | 12587010 | 1X |
| Primocin | Invivogen | ant-pm-2 | 100 µg/mL |
| L-glutamine | Sigma-Aldrich | G7513 | 2 mM |
| Human noggin | Peprotech | 120-10C | 100 ng/mL |
| Human EGF | Peprotech | AF-100-15 | 50 ng/mL |
| Human HGF | Peprotech | 100-39 | 50 ng/mL |
| Human FGF10 | Peprotech | 100-26 | 100 ng/mL |
| Human R-spondin-1 | Peprotech | 120-38 | 500 ng/mL |
| N-acetyl-L-cysteine | Sigma-Aldrich | A9165 | 1.25 mM |
| Nicotinamide | Sigma-Aldrich | 128275000 | 10 nM |
| A83-01 ALK inhibitor | Biotechne | 2939 | 500 nM |

**Table S3.** List of antibodies used in this study

| Protein | Manufacturer | Catalogue reference | Source, clonality, isotype | Assay | Dilution | Antigen retrieval method |
|---|---|---|---|---|---|---|
| Calponin | Millipore | ABT129 | Rabbit polyclonal | IF | 1 in 2,000 | Not applicable |
| CD10 | Santa Cruz Biotechnology | SC-46656 | Mouse monoclonal IgG1 | IHC | 1 in 1,000 | Citrate[1] pH 6.0, 2-3 h at 80°C |
| Collagen IV | Developmental Studies Hybridoma Bank (DSHB), University of IOWA | M3F7 | Mouse monoclonal IgG1 | IHC | 1 in 25 | Proteinase K[2], 20 min at RT |
| Cleaved Caspase-3 | Cell Signaling technology | 9664 | Rabbit monoclonal IgG | IF | 1 in 500 | Citrate pH 6.0, 2-3 h at 80°C |
| Fibroblast Specific Protein (FSP)-1/S100A4 | Abcam | ab218512 | Mouse monoclonal IgG1 | IHC | 1 in 8,000 | Citrate pH 6.0, 2-3 h at 80°C |
| pan-Cytokeratin | Sigma-Aldrich | c2562 | Mouse monoclonal IgG1/IgG2A | IF | 1 in 10,000 | Not applicable |
| | | | | IHC | 1 in 10,000 | Citrate pH 6.0, 2-3 h at 80°C |
| Podoplanin | NeoBiotechnologies | 10630-MSM1-P1ABX | Mouse monoclonal IgG1 | IHC | 1 in 15,000 | Citrate pH 6.0, 2-3 h at 80°C |
| Alpha-Smooth Muscle Actin (αSMA) | Sigma-Aldrich | a2547 | Mouse monoclonal IgG2a | IHC | 1 in 2,000 | Citrate pH 6.0, 2-3 h at 80°C |
| | | | | IF | 1 in 500 | Not applicable |
| Vimentin | Abcam | ab8978 | Mouse monoclonal IgG1 | IF | 1 in 800 | Not applicable |
| Vimentin | Abcam | ab45939 | Rabbit polyclonal IgG | IF | 1 in 400 | Not applicable |

[1]10 mM sodium citrate buffer; [2]20 µg/mL Proteinase K (Qiagen, 1014023).

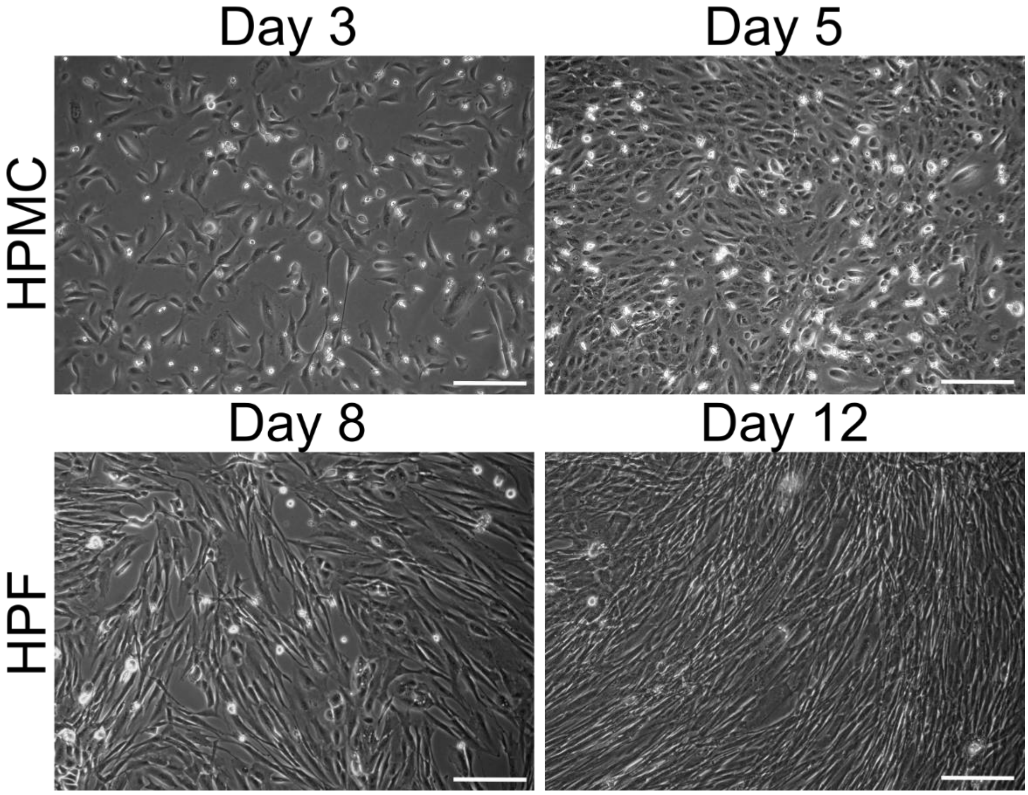


**Figure S1.** Human peritoneal mesothelial cells (HPMCs) and human peritoneal fibroblasts (HPFs) cell culture at passage 1 (P1), initial seeding density $1.67 \times 10^5$ cells/cm$^{2.}$ Scale bar: 200 μm

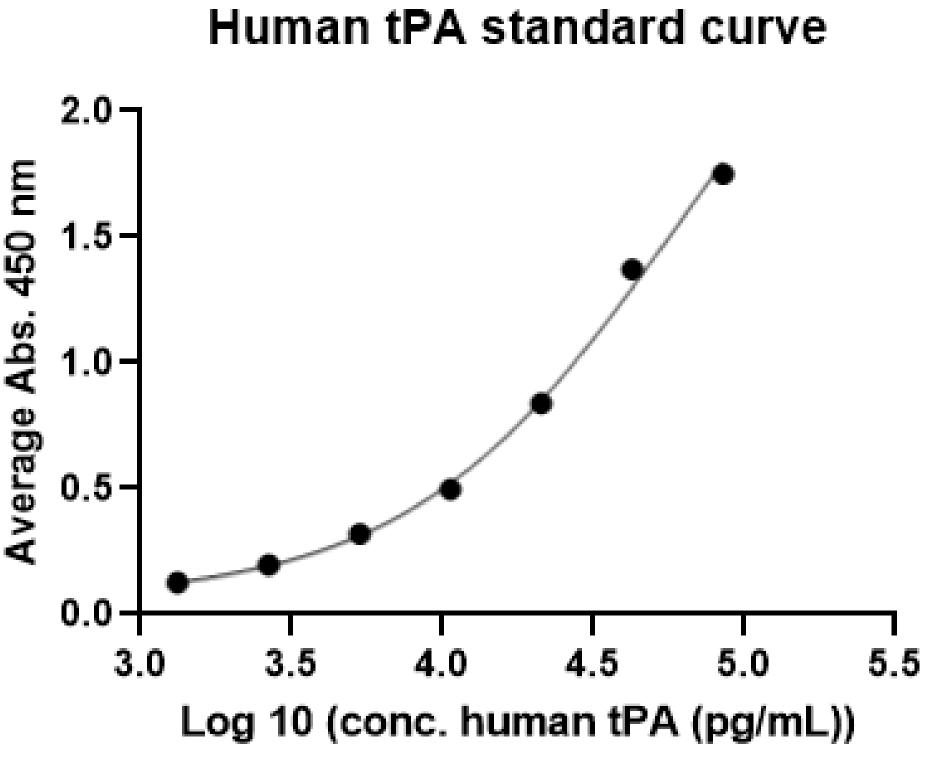


**Figure S2.** Standard curve for human tissue plasminogen activator enzyme-linked immunosorbent assay (tPA ELISA).

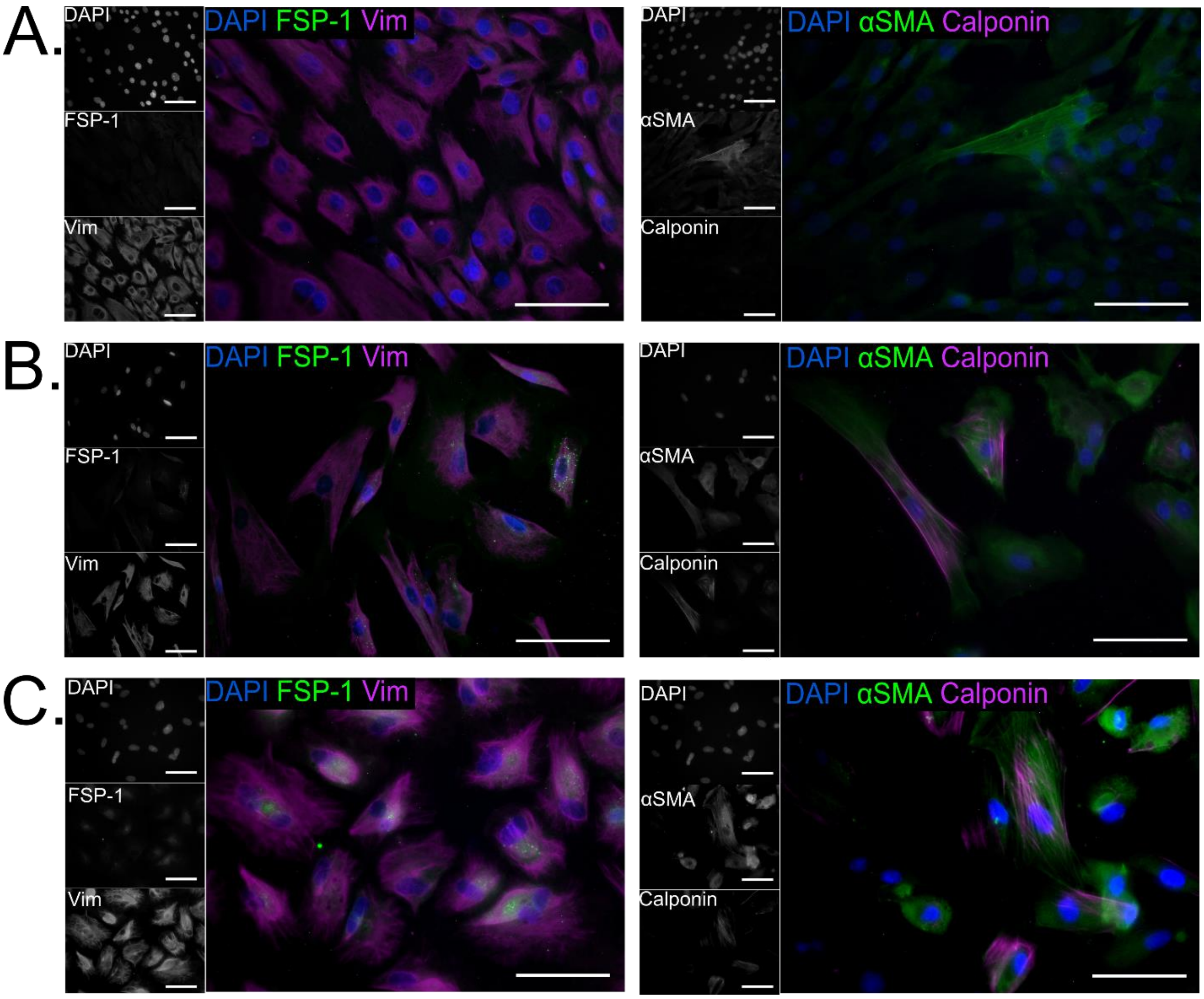


**Figure S3.** LP-9 cells at passage 10 (P10) under different culture conditions showing variability in fibroblast specific protein-1 (FSP-1), alpha smooth muscle actin (αSMA), and calponin expression. LP-9 cells were cultured under (A) mesothelial culture media (Medium199/MCBD105 (1:1 [v/v]) supplemented with 15% fetal bovine serum (FBS) and 10 ng/ml human epidermal growth factor (EGF)); (B) fibroblast growth medium (Promocell, C-23110) containing 1 ng/ml human basic Fibroblast Growth Factor (bFGF) and 5 µg/ml recombinant human Insulin; (C) Endometrial stromal cell culture medium (DMEM/Ham's F-12 media supplemented with 10% FBS). Scale bar: 50 µm.
